# ELF3 controls trait heterogeneity by tuning the rate of maturation in Arabidopsis and barley

**DOI:** 10.1101/2025.03.24.644900

**Authors:** Kayla McCarthy, Hannah Pook, Ethan Redmond, James Ronald, Joseph Keir, Zihao Zhu, Marcel Quint, Phillip A. Davis, Sue E. Hartley, Seth J. Davis, Daphne Ezer

**Affiliations:** Department of Biology, University of York, Wentworth Way, York YO10 5DD, United Kingdom; Institute of Agricultural and Nutritional Sciences, Martin Luther University Halle-Wittenberg, Halle (Saale), Germany; Leibniz Institute of Plant Genetics and Crop Plant Research, D-06466 Seeland, OT Gatersleben, Germany; Stockbridge Technology Centre, Stockbridge House, Selby YO8 3TZ, United Kingdom; Fischer Farms, Blaser Mills Law, 40 Oxford Road, High Wycombe, Buckinghamshire, HP11 2EE; School of Biosciences, University of Sheffield, Western Bank, Sheffield S10 2TN, United Kingdom

## Abstract

Trait heterogeneity in a population increases the likelihood that some individuals will survive an unpredictable environmental stress. Individual plants mature at different rates due to genetics, environmental factors, and random chance. All of this contributes to trait heterogeneity. *EARLY FLOWERING3 (ELF3),* a core circadian clock component, determines developmental timing, and here we tested the hypothesis that it would shape trait heterogeneity. We developed a model predicting that faster-developing populations exhibit greater heterogeneity during development but reduced heterogeneity at maturity, while slower-growing populations show the opposite pattern. Experiments in Arabidopsis *elf3* and barley *Hvelf3* mutants supported this prediction. In Arabidopsis, *ELF3* controlled hypocotyl elongation variability via maturation rate rather than direct regulation. Bolting time heterogeneity also decreased in faster-growing plants. In barley, *Hvelf3* altered growth heterogeneity, but this was initially masked by germination timing differences. Finally, smaller barley plants were found to be more resilient to osmotic stress, suggesting that *ELF3*-driven trait heterogeneity may contribute to bet hedging. These findings highlight how modifying developmental rates influences population-level trait distributions and stress resilience.

## Introduction

Biological variation in traits that cannot be accounted for by genetics or environmental conditions is often referred to as developmental instability (DI) (Klingenberg 2019) or as intrinsic noise (Eling et al. 2019). The presence of DI was initially controversial, because it is likely that some of the unexplained variability in traits is caused by unobserved variability in environmental conditions, such as the presence of microenvironments (Palmer 1996). However, it is increasingly clear that it is possible to control the degree of DI by modifying gene networks (Thattai and van Oudenaarden 2001), suggesting that the degree of trait variation is under evolutionary selection.

Phenotypic heterogeneity in a population can help make it resilient to unpredictable environments. Diversified bet hedging is the evolutionary strategy of having some individuals in a population have maladaptive phenotypes to the current conditions to maximise fitness over longer time scales (Simons 2009). For instance, a subset of a bacterial population will be in a persister state, in which reproduction is slowed but in which antibiotic resistance is improved (Verstraeten et al. 2015) Another classic example of diversified bet-hedging is heterogeneity in seed germination (Simons and Johnston 2006; Abley et al. 2021) and in egg hatching times (García-Roger et al. 2014), which make the populations more resilient to sudden environmental fluctuations. Because organisms will have different starting points in their development, they will be at different developmental stages during an extreme weather event increasing the chance that at least some individuals will be able to survive and reproduce.

An alternative way of diversifying the developmental phases within a population is by having variability in the rate of maturation (Simons and Johnston 1997); however, this has not been fully explored in the literature. Although initially proposed in the 1990s, there are few concrete examples linking heterogeneity in maturation rate to a bet-hedging strategy, with only one direct case reported (de Jong et al. 2010) and another from a simulation study on bet-hedging over developmental time (de Groot et al. 2023). The rate of maturation is a trait that falls under the umbrella term ‘heterochrony’, which refers to differences in the timings of developmental trajectories, between species, organisms, or different traits within the same organism.

In plants, there are close links between how plants regulate their maturation rates and how they maintain their daily circadian rhythms. The plant circadian clock is a complex gene network that confers a 24-hour period of oscillation to over a third of the Arabidopsis transcriptome (McClung 2019). The circadian clock is involved with detecting the duration of daylight in each 24-hour period (photoperiod) and so may help plants time their development appropriately with the seasons (Nomoto et al. 2013; Sanchez and Kay 2016). Almost every gene in the plant circadian clock has also been associated with the control of flowering time. For instance, EARLY FLOWERING3 (ELF3) is a core element of the Evening Complex (EC) of the plant circadian clock, but it was initially discovered because *elf3* flowers extremely early under non-inductive conditions (Zagotta et al. 1996; Ezer et al. 2017). Arabidopsis *elf3* has also been implicated in rapid hypocotyl elongation (Reed et al. 2000; Lu et al. 2012), early senescence (Sakuraba et al. 2014) and as an essential contributor to thermomorphogenesis and photomorphogenesis (Anwer et al. 2020; Zhu et al. 2022). In barley, *HvELF3* is also referred to as *EARLY MATURITY 8* (*Eam8*) and natural loss-of-function variants also result in more rapid development (Faure et al. 2012).

In addition to controlling developmental timing, *ELF3* sits in a quantitative trait locus controlling global gene expression heterogeneity (Jimenez-Gomez et al. 2011). Modifying the rate of maturation of a plant has the potential to influence the heterogeneity of traits, as has been seen with the Arabidopsis clock gene *ELF4* (Doyle et al. 2002), which forms a complex with ELF3 (Herrero et al. 2012). In this manuscript, we hypothesise that *ELF3* regulates trait heterogeneity indirectly, by modifying the plant’s rate of maturation. We began by developing a mathematical model that describes our hypothesis, predicting the properties of trait heterogeneity that we would have expected if *ELF3* indirectly modified trait heterogeneity by controlling the maturation rate of the plant. Then, we show that this null model generates predictions that are consistent with observations in Arabidopsis and barley, suggesting that *ELF3* indirectly controls trait heterogeneity. Finally, we demonstrate that *ELF3* can drive robustness to an environmental stress, specifically osmotic stress—a model of drought stress, by influencing the rate of maturation in barley. These results support that modifying the rate of maturation influences the spread of physiological traits within a plant population and may impact its robustness to environmental stresses.

## Results

### Variability in maturation rates lead to a trade-off between heterogeneity during growth or at maturity

Our aim was to determine whether *ELF3* modifies trait heterogeneity by adjusting the rate of maturation. To do this, we first needed to understand how adjusting the rate of maturation would influence trait heterogeneity. We did this by developing a mathematical model of plant development to generate predictions, which we later tested experimentally. To explore how maturation rate influences trait heterogeneity, we constructed a stochastic simulation of developmental progression. In our model, we assumed that each individual plant had its own internal *biological age* which could vary from its *chronological age* (time since seed was sown). Biological age was considered a latent variable, meaning that it can’t be directly measured. Measurable physiological traits were associated to a combination of both kinds of age, because traits related to growth are controlled by the developmental programme (i.e. the biological age) but may also be bottlenecked by accumulation over time in the plant (i.e. the chronological age)—for instance, plants may have a limit as to how quickly they can grow or produce leaves (Klingenberg 2019). Our model lets us tune the relative importance of biological age and chronological age in determining a physiological trait. In our model, the *biological age* completely determined the timing of key developmental transitions, such as the transition from vegetative to reproductive growth. Many physiological traits – like leaf size – have mostly stationary values after the vegetative-to-reproductive transition (Woo et al. 2016).

The scenarios that we modelled are summarised in **Table 1**. We modelled two underlying reasons why biological age varied from chronological age: (i) starting point variation – i.e. heterogeneity in germination times – and (ii) maturation rate variation – i.e. heterogeneity in how quickly plants age. Starting point heterogeneity has been extensively investigated as a bet-hedging mechanism (Simons and Johnston 2006; García-Roger et al. 2014; Abley et al. 2021), but maturation rate heterogeneity is understudied (Simons and Johnston 1997). The stochastic process of aging was simulated as a Gamma Process with drift (Lewis et al. 1989), which has previously been used to model biological ageing (Link and Hesed 2015).

**Table 1:**
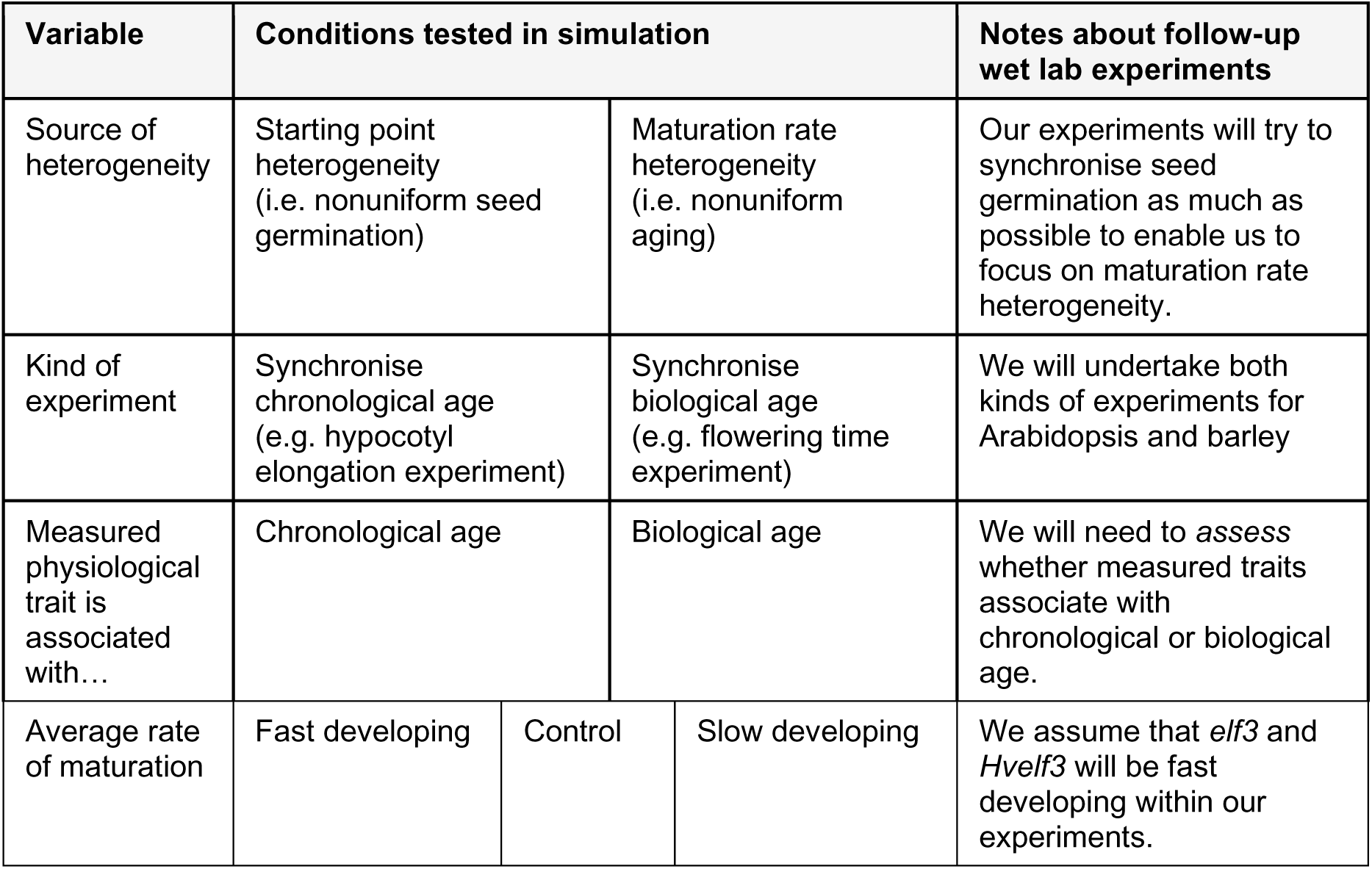
Summary of conditions tested in growth simulation.

We simulated biological age over time, under different average maturation rates (without changing the standard deviation of maturation rates) to reflect how we expected *ELF3* to influence the average maturation rate. We were also curious how starting point variation or maturation rate variation would modify trait heterogeneity. For this, we simulated two alternative scenarios: (i) where the only source of variation in biological age came from starting point variation. This is equivalent to saying that the plants would each germinate at slightly different times but then mature at the same rate (**Fig 1A**). (ii) where the only source of variation was maturation rate variation. This was equivalent to saying that the plants germinate synchronously but then have different rates of development (**Fig 1B**). In the first scenario, heterogeneity in biological age did not increase over time, because all plants developed at the same rate. In the second scenario, heterogeneity in traits increased over time, because faster maturing plants diverged from slower maturing plants over chronological time. Our modelling framework is flexible enough to enable future research into combinations of these two sources of heterogeneity.

**Figure 1:**
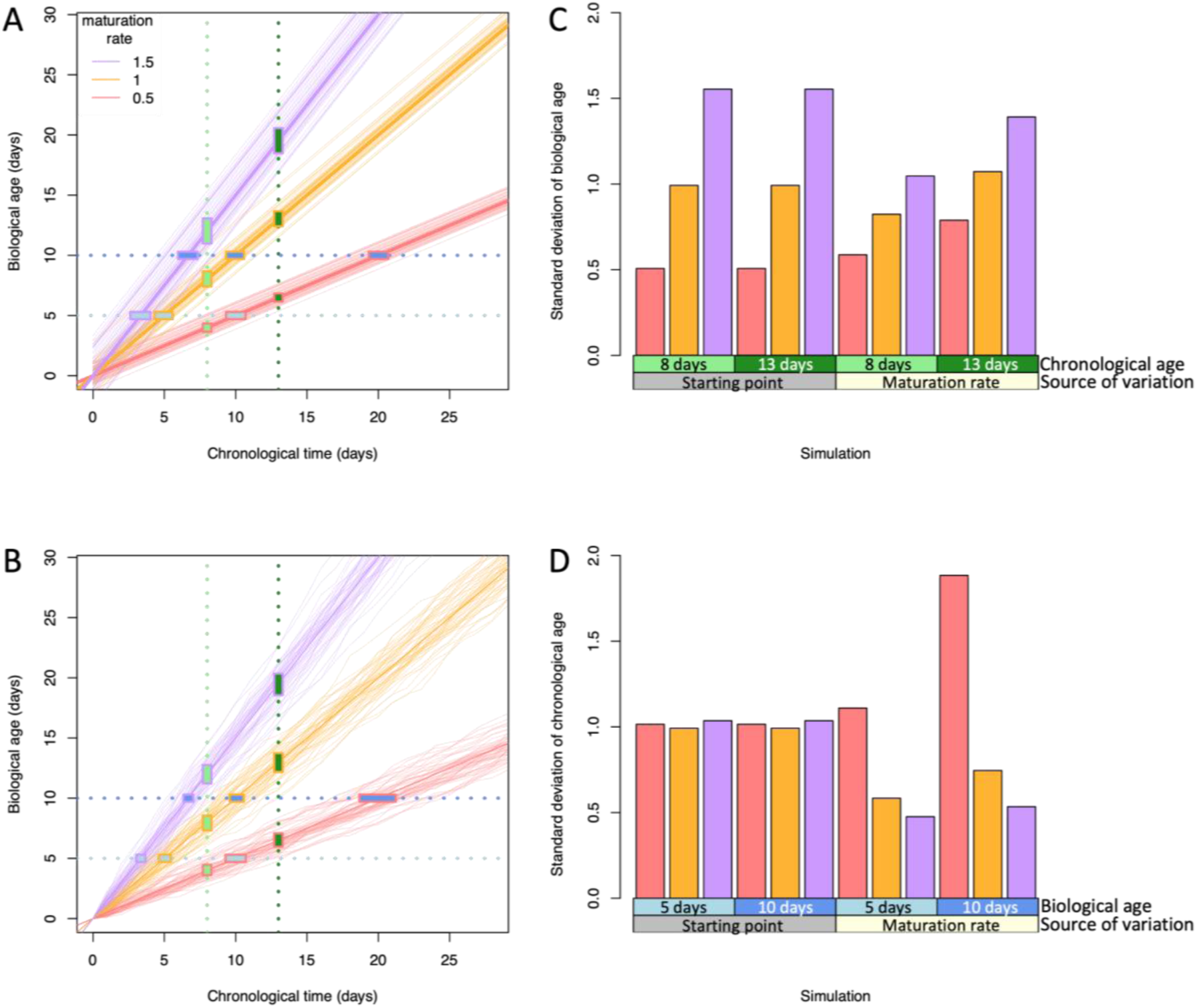
Maturation rate controls a trade-off in heterogeneity during growth and maturity. We generated simulations of heterogeneity (A) in starting point times, e.g. germination or egg hatching, where the starting points are drawn from a normal distribution (B) in maturation rates, where the growth curves are generated by a Gamma Process. In both cases, we consider three different average rates of maturation (purple=fast, orange=medium, pink=slow). Vertical lines highlight the situation in which plants are sampled at a specific chronological time, while horizontal lines highlight the situation in which plants are sampled once they pass a certain developmental transition. The rectangles indicate the interquartile range of the simulations under each of these experimental sampling schemes. The colours of these vertical and horizontal lines are made to be consistent with the boxes on the x-axis of figures C and D. We assume that some traits are associated with biological age, some with chronological age, and some with a mixture. Fig C highlights the amount of trait heterogeneity (standard deviation of biological age) when plants are sampled at a specific chronological time. Fig D highlights the amount of trait heterogeneity (standard deviation in chronological time) when plants are sampled at a specific biological time. Having a faster maturation leads to a lower standard deviation in chronological time at a specific biological age, but only in the presence of maturation rate heterogeneity. The colours of the lines and rectangles in (C,D) are consistent with the colour-code in (A,B).

Next, we wished to demonstrate how these theoretical results in **Fig 1A and B** would impact the measurements taken during two common plant physiology experiments. In the first kind of experiment, whose results are depicted in **Fig 1C**, we considered the experimental design in which a plant biologist would sow seeds, wait a certain number of days, and then measure a developmental trait. This was similar to the experimental design of hypocotyl elongation experiments (Ronald et al. 2024). Our results suggested that faster developing plants (such as in an *elf3* mutant line) always had greater trait heterogeneity. However, there were also qualitative differences in trait heterogeneity that differed in our two simulation scenarios. If there was heterogeneity in maturation rates (scenario 2), then the amount of variation increased over time.

In the second simulated experiment, whose results are shown in **Fig 1D**, we considered an alternative experimental design in which a plant biologist would phenotype their plants at a specific developmental stage, such as the onset of the vegetative-to-reproductive transition, (Redmond et al. 2024), in order to ensure that the plants are measured at the same biological age. In this case, we considered a trait that is associated with the chronological age of the plant, such as a trait that increased over time regardless of the biological age. If the only source of variation in biological age came from germination time, then modifying the average maturation rate would have no effect on trait heterogeneity. On the other hand, if there was heterogeneity in maturation rates, then faster developing plants had less trait variation than slower developing plants.

Given that many growth processes stall after the vegetative-to-reproductive transition (Woo et al. 2016), our model predicted that there is a trade-off between having trait heterogeneity during growth and at maturity. One might expect that plant populations that are more heterogeneous would continue to be more heterogeneous over developmental time, but our simulation suggested that this is not always the case. Instead, our simulation suggested that faster developing plants would have more trait heterogeneity during growth, but less trait heterogeneity at maturity, and vice versa. Since *ELF3* modifies the average rate of maturation (Zagotta et al. 1996), we expected to see this trade-off in any plant population that exhibits some amount of heterogeneity in the rate of maturation. This gave us testable model predictions, see **Table 2**.

**Table 2:** Summary of model predictions that we test experimentally.

| Experiment type | Trait(s) measured | If this assumption is met... | ...then, this is our model prediction. |
| --- | --- | --- | --- |
| Synchronised chronological age | Hypocotyl length (Arabidopsis)<br>Biomass (barley) | Traits are associated with biological age | Faster developing lines like *elf3* and *Hvelf3* will have **more heterogeneity** than controls. |
| Synchronised biological age | Leaf count (Arabidopsis)<br>Shoot fresh weight, leaf count, head count, tiller count (barley) | (i) There is some amount of maturation rate heterogeneity in the population (ii) Traits are associated with chronological time. | Faster developing lines like *elf3* and *Hvelf3* will have **less heterogeneity** than controls. |

### ELF3 may influence growth phase heterogeneity by adjusting the rate of maturation in Arabidopsis (Synchronised chronological age)

Maturation-rate variation can drive differences in trait heterogeneity, and *ELF3* may contribute to this by accelerating development. In the first simulated experiment (**Fig 1C**), we showed that increasing the mean maturation rate should increase trait heterogeneity in growing plants. We hypothesized that *elf3* has a faster maturation rate and would therefore lead to greater heterogeneity in hypocotyl lengths. Our previous work measured hypocotyl growth curves in Ws-2 wild-type and *elf3* Arabidopsis seedlings that could be used to test this prediction (Ronald et al. 2024). *ELF3* is involved in photoperiod detection (PMID: 31694066), so the experiment was performed under two different photoperiods, short day (SD, 8 hours light/16 hours dark) and long day (LD, 16 hours light/8 hours dark). This data confirmed that there was greater heterogeneity in hypocotyl lengths in *elf3* than in wild type Ws-2, as we expected (**Fig 2A, 2B**). The data also showed that *elf3* has greater heterogeneity in hypocotyl lengths than Ws-2 under both photoperiods. There was significantly lower heterogeneity of final hypocotyl lengths in Ws-2 than *elf3* under both photoperiods (p<0.05 in both cases, one-sided F-test for difference in variances, Holm’s adjusted).

**Figure 2:**
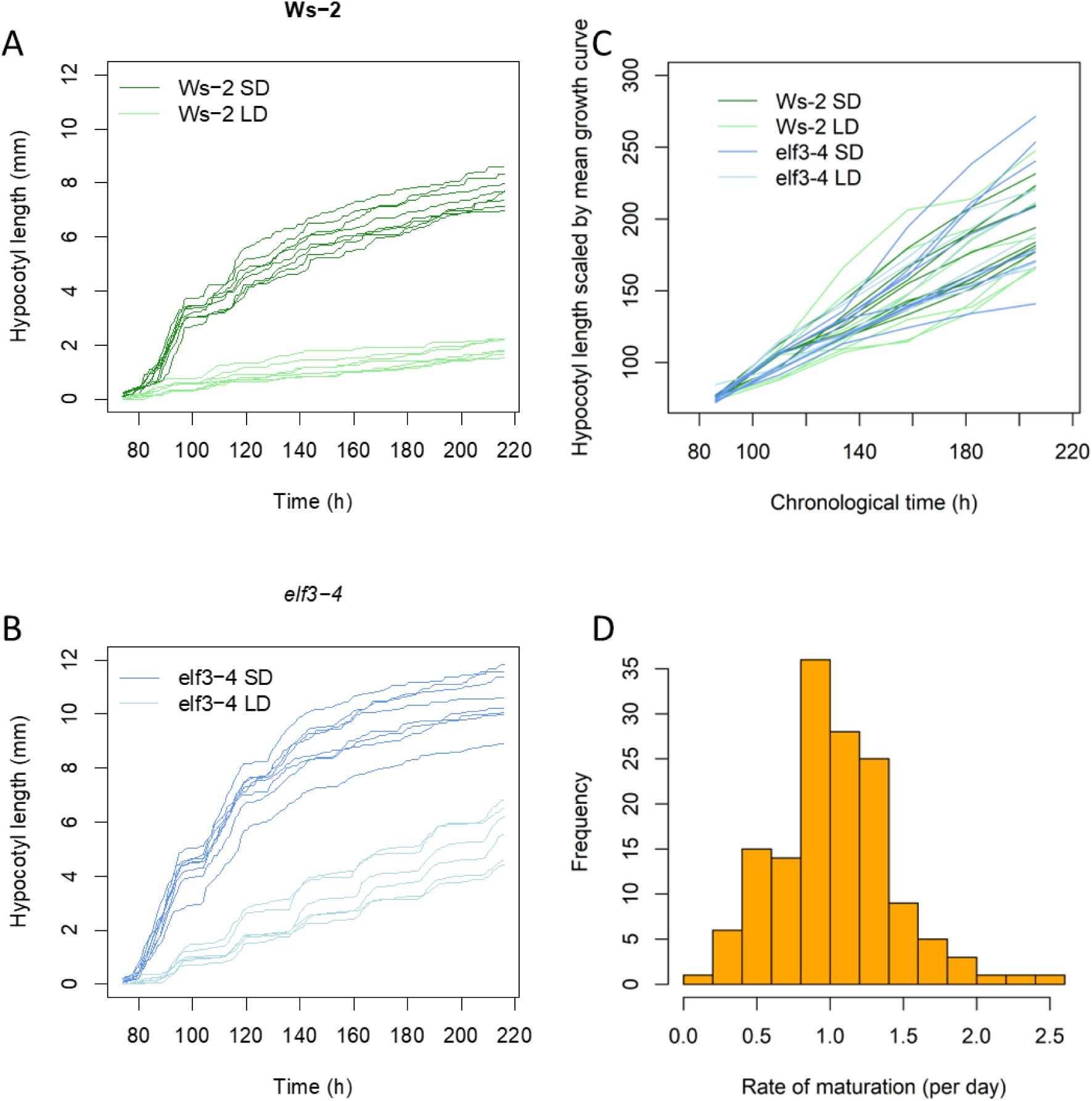
ELF3 influences trait heterogeneity during hypocotyl elongation by adjusting the average rate of growth. (A) Hypocotyl elongation was measured hourly using an infrared camera in Ws-2 (A) and elf3-4 (B) under short day (SD, 8hr/16hr light/dark cycles) and long day (LD, 16hr/8hr light/dark cycles). The height of each individual plant is drawn. (C) We calculated the average growth curve for each condition (genotype/light condition) and scaled each individual growth curve in reference to this mean curve. (D) The distribution of the daily growth rate of all the scaled individual curves in (C) is shown here.

To assess whether *ELF3* influences trait heterogeneity through maturation rate alone, one must examine if changes in mean maturation rate explain the observed variation in hypocotyl length. By testing this relationship, we can determine whether *ELF3’*s effect on trait heterogeneity arises solely from its influence on maturation rate or if additional regulatory mechanisms are involved.

To determine whether adjusting maturation rate is sufficient for explaining the changes in hypocotyl heterogeneity, we needed to carefully scale the growth curves of each plant by the average growth curve for their genotype and treatment group, in a way that would allow us to directly and fairly compare the plant’s degree of heterogeneity between genotypes and treatment groups– see **Fig 2C**. We scaled the curves to compare where each individual plant sits on its growth trajectory compared to the average plant in the population (its genotype/treatment group). If *ELF3* only adjusts the mean growth rate and not the spread in growth rates, then we would expect that *elf3* would have the same distribution of curves as Ws-2. Indeed, there were no significant differences between the scaled curves between genotypes or conditions (p>0.05, Kolmogorov-Smirnov test). The differences in trait heterogeneity observed between *elf3* and Ws-2 are sufficiently explained by *ELF3* control over the rate of maturation. Additionally, there is a fanning pattern in **Fig 2C** which is similar to what our simulation indicated in **Fig 1B**, which suggests that there is some heterogeneity in maturation rates.

Our simulation included as a core assumption that the distribution of maturation rates each day followed a gamma distribution, since we modelled growth as a gamma process. The analysis in **Fig 2C** can help us test whether this was an appropriate assumption. In **Fig 2D** we show the distribution of slopes in **Fig 2C**, which we would expect would follow a gamma distribution if hypocotyl length was associated with biological age. None of the four conditions had maturation rates that differed significantly from the gamma distribution based on the variance-ratio test (Villaseñor and González-Estrada 2015), and neither did the merged set of maturation rates significantly differ from a gamma distribution. This supports our use of a gamma process to model biological age over chronological time.

In summary, our data support the principle that *ELF3* modulates the heterogeneity in hypocotyl elongation by adapting the plant’s rate of maturation. Moreover, our data suggest that maturation rate follows a gamma distribution and that there exists some heterogeneity in the rates of maturation between individual plants.

### *ELF3* may influence heterogeneity during bolting by adjusting the rate of maturation in Arabidopsis (Synchronised biological age)

Our simulation suggested a trade-off between trait heterogeneity during growth and at maturity. Next, we wished to establish whether faster developing plants had lower heterogeneity during the vegetative-to-reproductive transition, as our model predicted.

However, our model predicted that this would only happen if two criteria were met: (i) there existed some amount of heterogeneity in maturation rates within the plant population, (ii) the trait was at least partially associated with chronological age, not just biological age. The fanning pattern in **Fig 2C** suggested that criteria (i) was met. To determine whether criteria (ii) was met, we sought to diversify the flowering time of Ws-2 plants by growing them in a wide range of photoperiods and then assessing whether leaf count was associated with flowering time (**Fig S1**). Our analysis confirmed that this second criteria was also met: leaf count was associated with chronological time.

To evaluate our hypothesis that faster developing plants have lower heterogeneity at bolting, we measured bolting time and leaf count at bolting in Ws-2 and *elf3* under the two photoperiods (SD and LD). As expected, *elf3* plants had reduced flowering time under both SD and LD conditions (**Fig 3A**), and there was a positive association between mean flowering time and standard deviation (**Fig 3B**), which was consistent with our hypothesis. Similarly, *elf3* had reduced numbers of leaves (**Fig 3C**) and standard deviation (**Fig 3D**). This confirmed that the faster developing *elf3* had less heterogeneity than Ws-2 at bolting, as our model predicted. Taken together with our previous results, we observed that faster developing plants had greater trait heterogeneity during growth, while slower developing plants had greater trait heterogeneity at bolting, as predicted by our model.

**Figure 3:**
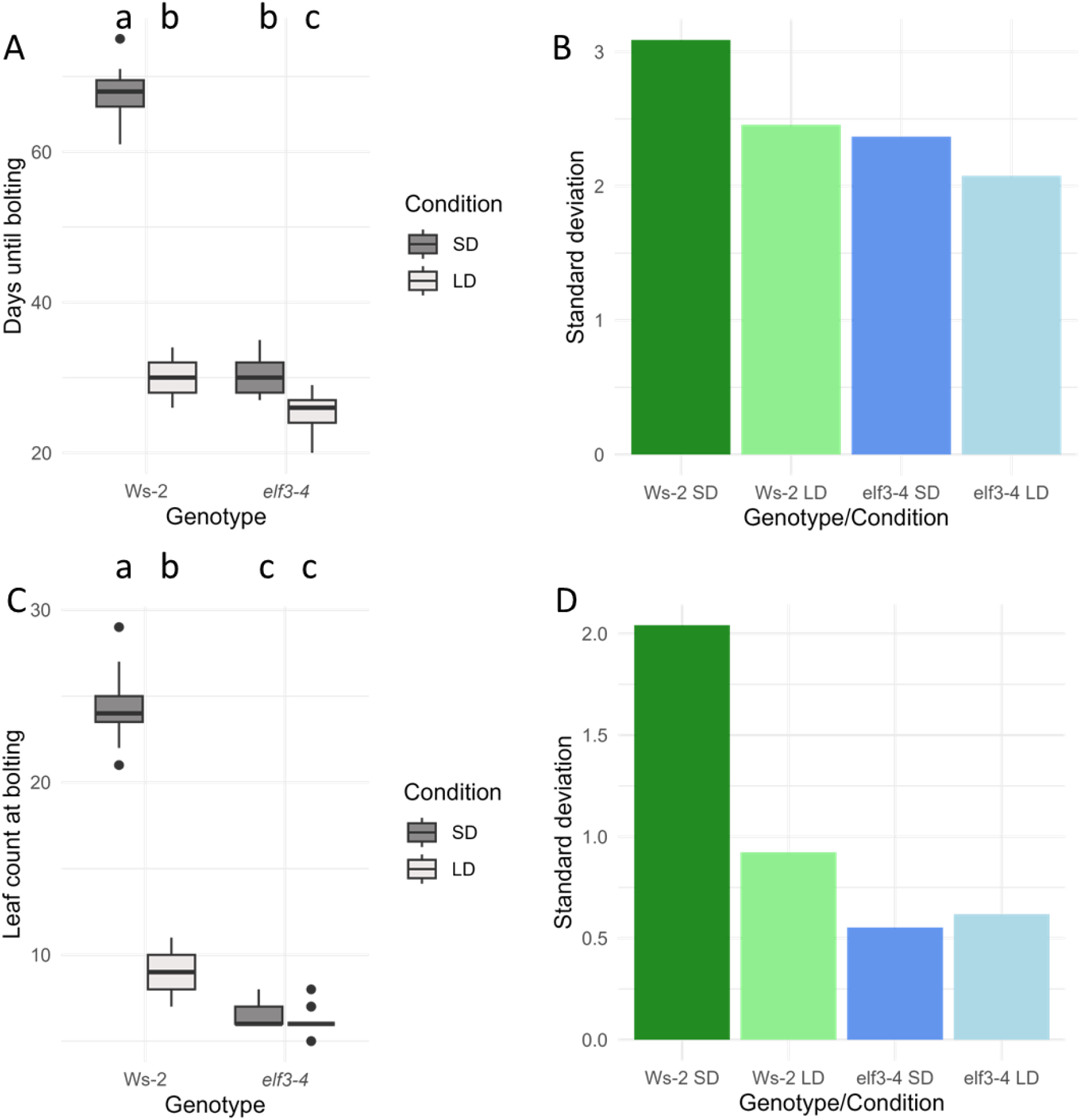
*ELF3*’s influence on adjusting maturation rate is sufficient to explain trends in trait heterogeneity at bolting. (A) Days to bolting under two photoperiods, short day (SD, 8hrs light/16 hrs dark) and long day (LD, 16hrs light/8 hrs dark) in Ws-2 and *elf3-4.* Box plots display medians, interquartile ranges, and the range excluding outliers. Outlier points are those that lie more than 1.5 times the interquartile range away from the median. Letters indicate groups that are statistically significantly different from one another, as defined by p<0.05 under a Tukey HSD test. (B) Standard deviations of the samples in (A) are shown. (C) Leaf count at bolting under two photoperiods, with standard deviations shown in (D).

### *Hvelf3* influences the germination and growth rate in barley (Synchronised chronological age)

Earlier in this study, we’ve shown that *ELF3* in Arabidopsis may influence the heterogeneity of phenotypes in a population in opposite directions, depending on whether the trait is measured at a consistent chronological time or at a specific developmental state. *Hvelf3* has similar early maturing properties as *elf3*, so we investigated whether the trade-off in Arabidopsis between growth and trait heterogeneity extended to barley, by investigating biomass accumulation in Bowman barley in two variants of *Hvelf3*, which we refer to as *eam8.k* and *eam8.w* (see Methods). We focussed on fresh biomass rather than a physiological feature more analogous to hypocotyl length because this was easier to measure, as barley plants have multiple tillers.

Before performing this experiment, we optimised the conditions to synchronise the germination as much as possible, observing that germination rates were saturated after 3 days of 4 °C exposure (**Fig S2**). However, there were still significant differences in germination times between genotypes. For this reason, we shifted the growth curves so the x-intercepts (i.e. germination times) were aligned at time point zero (**Fig S3**) to enable us to focus on the impact of *Hvelf3* on heterogeneity in maturation rates. The aligned growth curves are shown in **Fig 4A**. To enable a fair comparison between genotypes, we scaled the growth curves by the mean curve for each condition **Fig 4B**, producing a figure analogous to **Fig 2B**. Just as in Arabidopsis, we found that there was a fanning out of scaled growth curves that was consistent with heterogeneity in maturation rates, and there were no clear differences between genotypes after scaling.

**Figure 4:**
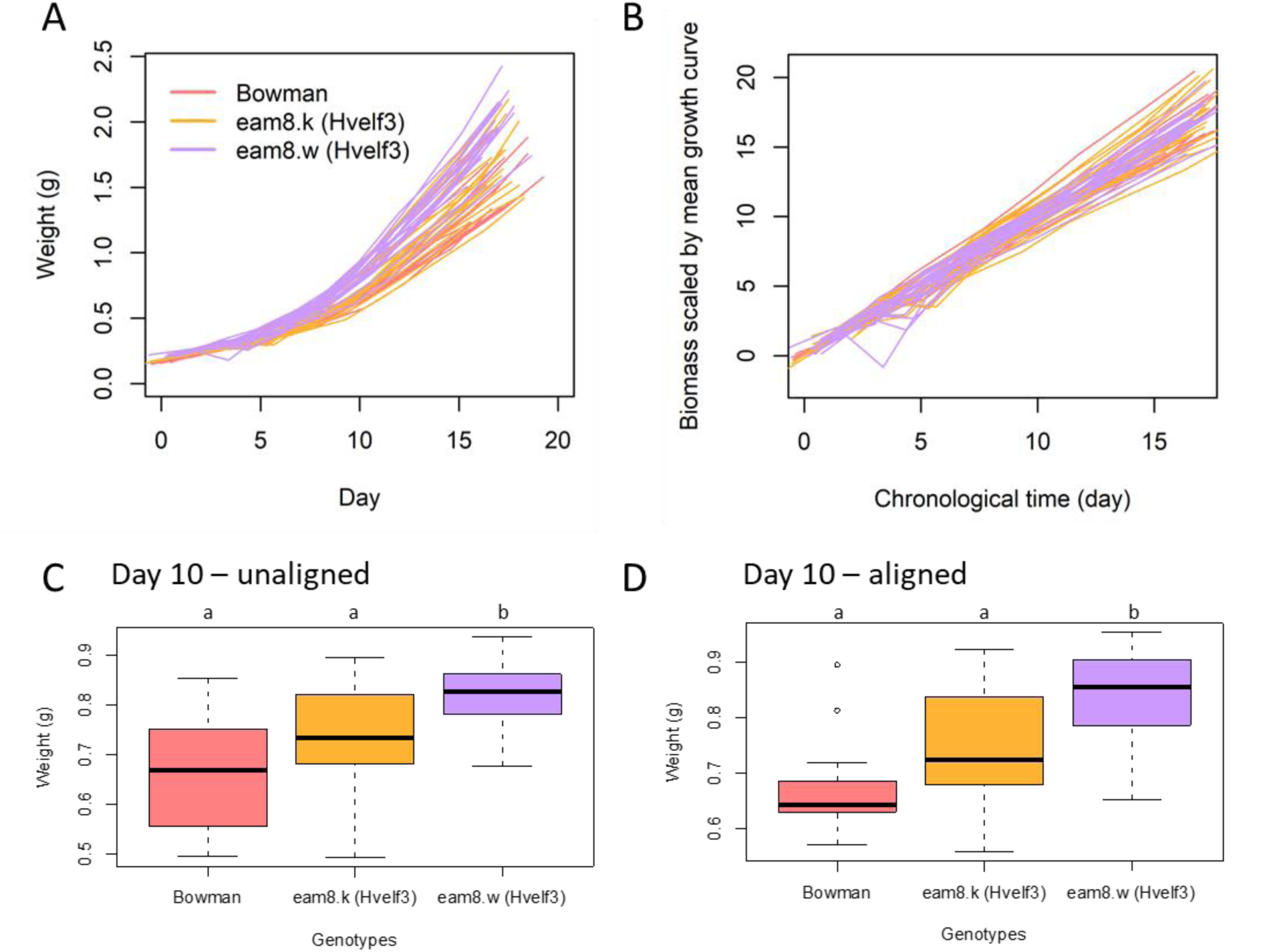
*Hvelf3* is associated with faster accumulation of biomass and more trait heterogeneity, but only once growth curves are aligned by germination time. (A) Fresh weight (g) measured in barley from the time point they were transferred to Hoaglands media. Each line represents an individual plant, colour-coded by genotype, where *eam8.k* and *eam8.w* have variants in *Hvelf3*. (B) These individual growth curves were then scaled by their mean growth curves to enable fair comparison between genotypes. Next a comparison of fresh weight (g) at a single chronological time point, day 10, was made before (C) and after (D) aligning their germination times. Letters indicate groups that are statistically significantly different from one another, as defined by p<0.05 under a Tukey HSD test. Box plots display medians, interquartile ranges, and the range excluding outliers. Outlier points are those that lie more than 1.5 times the interquartile range away from the median.

Bowman and *eam8.w* had significantly different means by day 10 (**Fig 4C**). Intriguingly, our data suggested that *Hvelf3*’s effect on barley germination and its effect on growth rate have opposing impacts on trait heterogeneity. Before we aligned the growth curves to synchronise germination times, the two *Hvelf3* genotypes *eam8.k* and *eam8.w* appeared to have less dispersion than Bowman (**Fig 4C**). However, once the growth curves were aligned by their germination times, both *Hvelf3* genotypes had greater heterogeneity in biomass (**Fig 4D**). This suggested that faster growth rates may be correlated with greater trait heterogeneity, but this may be masked by *Hvelf3*’s influence on germination time in barley.

### Barley trait heterogeneity at maturity differs for traits associated more with chronological time (Synchronised biological age)

Next, we wished to establish whether *Hvelf3* influences trait heterogeneity at the onset of flowering in barley. We included variants of other early flowering lines in barley in our experiment: *eam10.m*, which includes variants in the barley ortholog of *LUX ARRHYTHMO (LUX), Hvlux*, —another constituent of the evening complex with ELF3 (Helfer et al. 2011)--and *eam5.x*, which includes variants of the barley ortholog of the photoreceptor PHYTOCHROMEC (PhyC), *HvPhyc*. Our model predicted that there would be less heterogeneity in traits in these early flowering lines, but only if the traits were associated with chronological time and were not solely determined by the developmental stage of the plant. We measured four different traits at the onset of flowering: fresh shoot biomass (**Fig 5A**), leaf count (**Fig 5B**), head count (**Fig 5C**) and tiller count (**Fig 5D**), which we compared with flowering time. The first two of these variables appeared to more closely correlate with chronological age than the latter two traits (R=0.795 and 0.756 for biomass and leaf count respectively, and R=0.436 and 0.324 for head count and tiller count, respectively). Our model would suggest that there would be a positive relationship between mean and trait variability in the traits associated with chronological age, but not those associated with biological age. Indeed, we observed this to be the case in **Fig 5E-H**. The only exception to this pattern appeared to be one of the *Hvelf3* variants– *eam8.w*– which had more variation than would be expected. With the exception of the *eam8.w* line, our results confirmed that fast developing plants have less trait heterogeneity at maturity, suggesting that the rate of development may indirectly influence trait heterogeneity, in a way that is consistent with our initial model, although this perhaps cannot explain trait heterogeneity for *eam8.w*.

**Figure 5:**
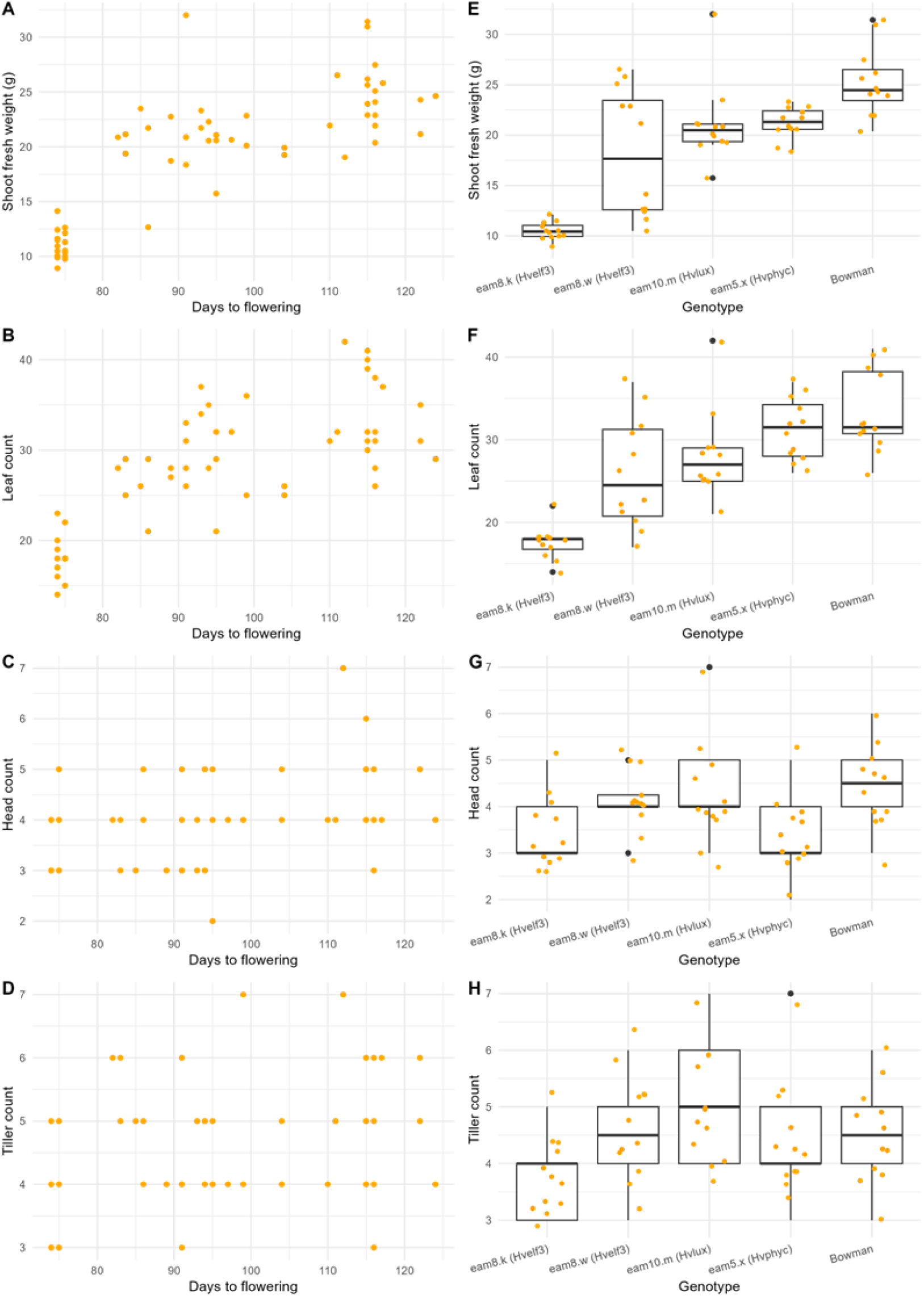
Heterogeneity of traits at maturation depend on association between chronological age or biological age. Shoot fresh weight, g (A), leaf count (B), head count (C) and tiller count (D) were measured at flowering time. We observe that traits that are more strongly associated with flowering time (A,C) have a greater association between mean and the interquartile range (B,D) with the exception of *eam8.w*, compared to traits that have a weaker association with flowering time (E-H). Box plots display medians, interquartile ranges, and the range excluding outliers. Outlier points are those that lie more than 1.5 times the interquartile range away from the median.

### *Hvelf3* and initial biomass impact response to osmotic stress in barley

We have suggested that *ELF3* was associated with developmental rate and therefore had an influence on trait heterogeneity during growth and at maturity. However, we wished to establish the fitness consequences of diversifying these traits.

First, we wished to determine whether the fresh biomass of a developing barley plant would influence its ability to withstand environmental stress. Our aim was to focus on a trait where we suspected that early maturing mutants would *not* directly impact stress response, but where the size of the plant *would* impact the response. Inducing osmotic stress in hydroponically-growth barley by introducing 15% polyethylene glycol (PEG) into the hydroponic media has previously been used as a model for drought response (Hellal et al. 2018). We performed an osmotic stress experiment on early maturing barley lines at a time point before the genotypes did not yet have significantly different biomasses. We found that the response to osmotic stress was associated with initial plant size, but not with genotype (**Fig S4**).

Therefore, we investigated how heterogeneity in the rate of growth would impact the capacity of barley to respond to osmotic stress. Our analysis in **Fig 4** demonstrated that *Hvelf3* may influence germination, but we wished to isolate the impact of maturation rate heterogeneity. To control for this factor, we measured when each individual plant surpassed a certain biomass threshold and then waited an additional 14 days before providing a polyethylene glycol (PEG) treatment to induce an osmotic stress. This would ensure that any heterogeneity in biomass was a result of the maturation rate heterogeneity, rather than starting point heterogeneity. In all three lines there was a negative association between pre-stress biomass and change in fresh weight (**Fig 6A**), with less biomass loss being associated with greater greenness of the barley. *Hvelf3* lines had a significant increase in biomass loss compared to Bowman after PEG treatment (**Fig 6B**). To determine whether differences in size distributions were sufficient to explain the differences in genotypes, we normalised the biomass loss by the pre-stress biomass (**Fig 6C**), and Bowman and *eam8.k* no longer showed statistically significant differences. This suggested that *Hvelf3* influenced osmotic stress response by regulating size, but this is not sufficient for explaining the results in all variant lines.

**Figure 6:**
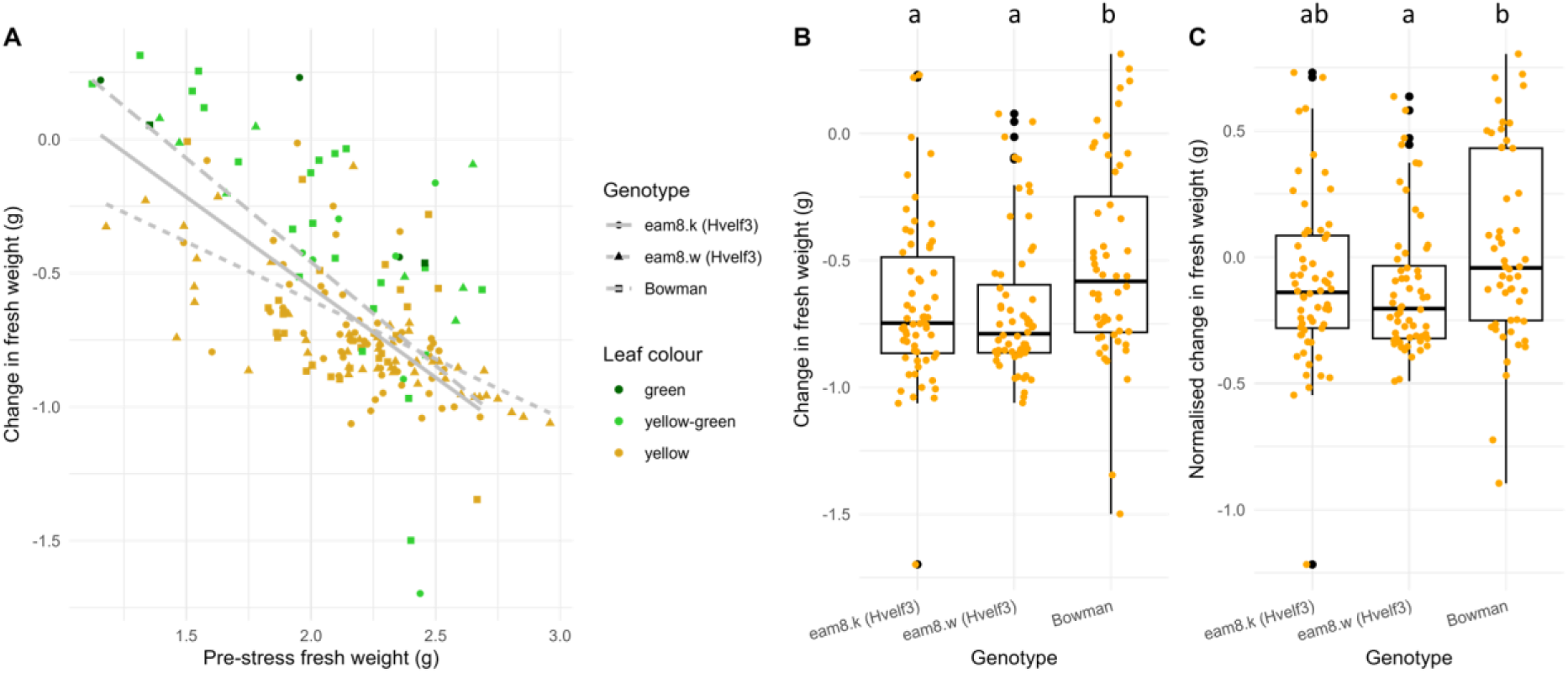
Impact of size on response to osmotic stress in barley. (A) Here we show that the initial fresh mass of the barley is negatively associated with a change in fresh mass after 15% polyethylene glycol (PEG) treatment in all lines (two *Hvelf3* variant lines and Bowman). Each plant is colour-coded by the leaf colour, indicating that change in fresh weight is associated with leaf yellowing. Lines of best fit are shown for each line. (B) *Hvelf3* variant lines show a larger magnitude change in fresh mass than Bowman. Box plots display medians, interquartile ranges, and the range excluding outliers. Outlier points are those that lie more than 1.5 times the interquartile range away from the median. Letters show statistically significantly different groups, as determined by p<0.05 in a Tukey HSD test. (C) A boxplot where the change in fresh mass was normalised by the prestress fresh weight.

## Discussion

### Maturation rate can lead to a trade-off between trait heterogeneity at growth and maturity: implications for life history evolution

In this study, we explored how *ELF3*—a circadian clock gene that controls developmental timing--can modulate trait heterogeneity in plant populations. We first built a mathematical model to assess how shifts in maturation rate, distinct from variation in initial germination, can drive diverse phenotypic outcomes. The model predicted a trade-off in trait dispersion during the growth phase versus at maturity: faster-growing populations accrue more variability early on but exhibit reduced variability when they reach a developmental transition. These predictions were then tested in Arabidopsis and barley lines with known mutations in *ELF3* or the barley *ELF3* homolog, *HvELF3*. Our work also reveals that barley biomass is associated with its capacity to withstand osmotic stress, suggesting that this heterogeneity may contribute to fitness under fluctuating environments.

Taken more broadly, this trade-off could drive the evolution of life histories of species (Tuljapurkar et al. 2009; Scheiner 2014). For instance, fast-developing organisms like dandelions have a lot of variability during growth due to being in the different developmental stages, but mature dandelions have less physiological variation than slow growing organisms like trees. Analysis of fish data suggests that heterogeneity of traits during growth and maturity may indicate evolution of life history strategies due to external stresses (Ma et al. 2021). Here, we propose that this same kind of trade-off is occurring, at a smaller scale, across genetically different sub-populations of the same species.

### Circadian clock via *ELF3* may contribute to this trade-off by controlling the maturation rate

Previous work has shown that the circadian clock is associated with gene expression noise. Studies have reported that *ELF3* is associated with global noise in gene expression (Jimenez-Gomez et al. 2011). ELF3 forms the evening complex with ELF4 and LUX, and ELF4 has also been implicated in controlling trait heterogeneity (Doyle et al. 2002). Moreover, ELF4 expression levels in seedlings can be used to predict the heterogeneity of flowering time in Arabidopsis (Lock et al. 2025). There is some evidence that the Evening Complex could directly regulate gene expression noise. For instance, the Evening Complex is associated with H2A.Z deposition (Tong et al. 2020) which is associated with circadian-dependent levels of gene expression noise (Cortijo et al. 2019).

However, the circadian clock can also indirectly impact trait heterogeneity by controlling the ageing process. Almost all circadian clock genes have an impact on flowering time (Sanchez and Kay 2016). Moreover, the robustness of circadian rhythms has been linked with life-spans across diverse organisms, including in humans (Acosta-Rodríguez et al. 2021; Cheng et al. 2022; Johnson et al. 2023). Here, we show that one of the implications of this is that by modifying developmental rates, the circadian clock can influence trait heterogeneity at different life stages within populations of organisms. Except for in the barley *Hvelf3* line *eam8.w,* all our results are consistent with *ELF3* having an indirect role in controlling trait heterogeneity. To say with confidence that a gene regulates trait heterogeneity, it is paramount that the study properly controls for developmental asynchrony.

### Maturation rate may be a contributor to diversified bet hedging strategies

Maturation rate heterogeneity may contribute to a diversified bet-hedging strategy. We showed that individual plants in a population will have different rates of maturation and that the amount of heterogeneity can be tuned by genetic variation. We also showed that the resulting differences in biomass in barley influences the amount of leaf senescence in response to osmotic stress. These properties are all consistent with maturation rate contributing to a diversified bet hedging strategy, but to further support this hypothesis, we require further research on the impact of developmental rate on fitness.

Diversified bet hedging requires that some offspring have maladaptive traits to the current conditions to enable greater robustness of the population to potential future adverse conditions (Abley et al. 2024). In Arabidopsis, it was found that there was selection pressure to flower early under most—but not all—sites tested (Fournier-Level et al. 2022). A previous meta-analysis has also revealed that selection for more rapid onset of flowering is widespread across plant species (Munguía-Rosas et al. 2011), with various potential mechanism proposed to explain this phenomenon (Austen et al. 2017). Taken together with our work, this suggests that slower developing plants may be less fit under optimal conditions but may have increased fitness under osmotic stress. However, faster development is not beneficial for fitness in all species and conditions. The adaptation of developmental rates is complex (Ehrlén 2015), and its integration into diversified bet hedging strategies warrants future investigation.

In summary, our research suggests that *ELF3*, a component of the plant circadian clock, may control the rate of development in a way that increases the amount of trait heterogeneity during growth and decreases trait heterogeneity at maturity. This suggests a novel mechanism by which the plant circadian clock could tune trait heterogeneity throughout the plant’s life cycle.

## Materials and methods

### Modelling starting point and maturation rate heterogeneity

We use a gamma process to model how biological age varies over chronological time. The user is required to provide *∈_A_* which influences the amount of heterogeneity in maturation rates, *∈_B_* which influences the amount of heterogeneity in starting points, and k which specifies the mean maturation rate.

The initial biological time *β_N_* is drawn from a normal distribution.

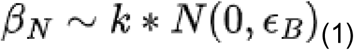

The change in biological age each day dX(t) is drawn from a gamma distribution, determined by a shape parameter alpha and scale parameter beta.

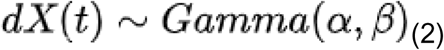

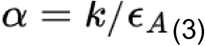

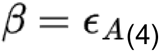

The choice of parameters in the gamma distribution ensures that the mean dX(t) is k and the variance is *∈_A_*.

There is strong biological motivation for using a gamma process and it has previously been used to model growth (Link and Hesed 2015). A gamma distribution models the waiting time for the Nth event to occur. The timing of biochemical events within an organism usually follow an exponential distribution, including protein binding, enzymatic reactions, etc and this has been the basis of the most common kind of stochastic simulation of biochemical processes (Gillespie 2007). Here we are essentially modelling ageing as a process that occurs each night until the number of active growth inhibitors exceeds a certain threshold. For instance, PHYTOCHROME INTERACTING FACTORs (PIFs) are a transcription factor that stimulate growth daily until active phytochromes inhibit them (Nomoto et al. 2013) In such a circumstance, a gamma distribution would describe the lengths of times until N inhibitors successfully bind the growth regulators.

A useful property of a gamma process (which makes it preferable to the more common ‘Gaussian process’) is that it ensures that maturation rate is always positive. However, it assumes that each day’s maturation is independent of the previous day’s rate, which is likely not the case, but is a useful simplification.

### Normalising growth curves by their average trajectory

We follow the schema of describing developmental instability which postulates that there is an idealised growth trajectory, but that noise, environment, genetics, etc causes variation from that idealised trajectory (Klingenberg 2019).

Let us call this idealised growth trajectory g(t), where t is time. For any individual plant, we can measure the actual growth over time as h_i(t) for the i-th plant in the population. Let us define b(t) as the biological time over chronological time.

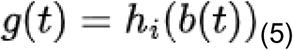

We can calculate biological age via:

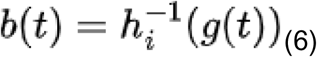

Now, we use this same formula to help us normalise each individual growth curve by the average growth curve for that population. We calculate g(t) as the mean growth curve of the population. We can calculate 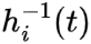 using the data that we collect about each individual plant’s growth data (hypocotyl length for Arabidopsis and biomass for barley). As growth is monotonic, we simply fit a separate model for each individual plant to predict its chronological age of the plant on the basis of its height or biomass, using smooth B-splines in the fda package in R (Ramsay and Silverman 2013). We use 200 B-splines and set the smoothing parameter lambda to 0.00001 for the Arabidopsis data and 8 B-splines and set the smoothing parameter to 0.0001 for barley. The growth curves in **Fig 2A,B** and **Fig 4A** also depict the fitted curves, using these same parameters.

### Arabidopsis experiments

Hypocotyl experimental data was previously published (Ronald et al. 2024) and consist of *elf3-4 CCR2::LUC* and Ws-2 *CCR2::LUC Arabidopsis thaliana* plants (Hicks et al. 2001; Ronald et al. 2022). In summary, sterilized Arabidopsis seeds were stratified (4°C, dark) for 72 hours on solid *A. thaliana* solution (ATS) nutrient medium (Lincoln et al. 1990) with 1% w/v sucrose. Vertical plates were kept at 21°C in constant light (LL, 40 μmol m-2s-1) for one day to induce uniform germination of seeds, before being to either long day (LD, 16 hrs light/8hrs dark) or short day (SD, 8hrs light/16hrs dark) conditions. Hypocotyl elongation measurements started at 72 hours after sowing (ZT0), by illuminating the plants by two infrared light emitting diodes and imaging them hourly by a Panasonic G5 camera modified with a long pass filter (830 nm). Image stacks were imported as “Image Sequence” in ImageJ, and the hypocotyl length was measured using the “Segmented Line” tool, by slightly adjusting the control nodes between time points.

For the flowering time experiment shown in **Fig S1**, Ws-2 *CCR2::LUC* seeds were stratified in ddH20 for 10 days at 4°C. After stratification, seeds were transferred to F2+S compost (P40 insert) supplemented with equal parts coir and vermiculite along with 0.7 g/L osmocote. Seeds were then transferred to a short-day (SD, 8hr light, 16 hr darkness) growth chamber with a constant temperature of 21°C for 21 days. On day 21, trays were transferred to a long-day (LD, 16 hr light, 8 hr darkness) chamber with a constant temperature of 21°C apart from one tray which stayed under SD (‘constant SD’ control). Light intensity under LD and SD was ∼100 µmol/m-2/s-1. Trays were then progressively moved back from the LD chamber to the SD chamber every day for one week apart from one tray which was kept under LD (‘constant LD’ control). Signs of flowering were then observed daily. Flowering was measured as the point at which the bolt was ∼1 cm above the height of the rosette. Days to bolting was also measured from the time stratification ended.

For the flowering experiments in Ws-2 (**Fig 3**), Ws-2 *LHY:LUC* (McWatters et al. 2007) and *elf3-4 LHY::LUC* (Herrero et al. 2012) seeds were surface sterilised and plated onto 1x MS plates (1.5% agar) supplemented with 0.25% w/v sucrose. After four days of stratification at 4°C, plates were transferred to a neutral day photoperiod (12h light/12h dark, 22°C) for 12 days. On day 12 days, seedlings were transferred to soil (1 part F2+S compost, 1 part coir and 1 part vermiculite supplemented with 0.7 g/L osmocote) and then moved to short-(8h light / 16 h darkness) or long-day (16h light / 8 h darkness) photoperiods with a constant temperature of 22°C and a light intensity of ∼100 µmol/m-2/s-1. Flowering was determined as the point at which the bolt was ∼1 cm above the rosette. The number of rosette leaves (including cotyledons) and days to flower since stratification ended was measured.

### Barley growth curves

The spring barley cultivar Bowman and introgression lines Bowman (*eam8.w/ BW290*) and Bowman (*eam.k/BW289*) both possessing the recessive *Hvelf3* allele, were used for this experiment (Faure et al. 2012; Zakhrabekova et al. 2012). Barley plants were grown hydroponically so that it would be possible to measure the fresh weight of individual plants over time. Please see the Appendix in the Supplementary Materials for further details of these lines.

Seeds were soaked in a solution of 33% bleach and 200 µL/L triton-X for up to 1 hour before being thrice rinsed in distilled water. Seeds were germinated on paper towel folded into 120/120/17 MM vented square petri dishes (Greiner BIO-ONE, Cat. No. 688102) with 50 mL distilled water, and then kept in the dark at 4 °C for 5 days.

Then, plates were moved to a controlled environment growth chambers– 210 µmol m-2 s-1 light intensity with 12 hour light/dark cycles and fluctuating temperatures 20°C light : 18°C dark with an ambient relative humidity (60% +/− 10%) throughout all barley experiments. Seeds were monitored daily for germination. The majority of seeds germinated within 48 hours of each other and only these seeds were moved forward.

Three days after the second germination date of the retained seeds, seedlings were transferred into 50 mL centrifuge tubes filled with a modified version of half strength Hoaglands solution (0.65M KNO3, 0.2M K2HPO4 · 3H2O, 0.4M Ca(NO3)2 · 4H2O, 0.2M MgSO4.7H2O, 0.46mM H3BO3, 0.05M MnCl2 · 4H2O, 0.2mM ZnSO4.7H2O, 0.1mM Na2MoO4, 0.2mM CuSO4, 0.45m MC10H12FeN2NaO8) (–0.2 MPa) as previously used (Habte et al. 2014) and secured to the top with a foam bung. Fresh weight of each individual plant was measured on the following days after being transferred: 1, 3, 5, 8, 12, 15, 17, and 19. This was done by removing the plant from the tube, gently drying the roots with a paper towel to remove excess water and then weighing their total mass, before placing the barley back into the tube.

### Barley flowering time experiments

This experiment used Bowman and the introgression lines Bowman (*eam8.w/ BW290*) and Bowmon (*eam.k/BW289*). In addition, we used Bowman (*eam10.m/BW289*) with the recessive *Hvlux* allele (Campoli et al. 2013), Bowman (*eam5.x/BW285*) which overexpresses *HvPHYC (eam5*) (Pankin et al. 2014). Please see the Appendix in the Supplementary Materials for further details of these lines.

The same seed sterilisation and stratification protocol was used, but barley was transferred to soil (1 part F2+S compost, 1 part coir and 1 part vermiculite supplemented with 0.7 g/L osmocote) and grown in a growth cabinet **(Snidjer,** 250-300µmol/m²/sec, 12-h light, 20 °C/12-h dark, 18 °C, LD treatments, ambient relative humidity 60% ± 10% RH) and monitored regularly until visible signs of flowering (GS49-51 as described in the Agriculture and Horticulture Development Board (AHDB), UK 2023 which aligns with the Zadoks decimal growth scale (Zadoks et al. 1974). Then, shoot fresh weight (g), leaf count, head count and tiller count were measured for each individual plant. These experiments were done in soil, rather than hydroponically, because our hydroponic set-up is too small to support mature barley plants. The end of booting (GS49) and beginning of ear emergence (GS51) was rather than flowering (GS61-69) because the Bowman cultivar is self-pollinated meaning their stigmas are receptive and capture sufficient self pollen before anther exertion (GS61). Flowering organs mature between awn emergence (GS49) and anther exertion while maturation of the anthers is usually under the sheath. Awn emergence (GS49) was chosen as it is the actual “flowering time” stage of spring barley as anthesis/fertilization happens around this stage. Sometimes the ear would emerge from the sheath (GS50-51) before the awns were visible (GS49).

### Barley stress experiment - early development

This experiment included all the same lines as the barley flowering time experiment as well as the Elite winter cultivar Antonella. The same seed sterilisation, stratification, and growth conditions were used as in the barley growth curve experiments.

Currently, drought stress models rely on presumably non-permeable high-molecular weight osmolyte polyethylene glycol (PEG) with an average molecular weight of 6000 Da or more (Hohl and Schopfer 1991). It is well documented that PEG effectively decreases medium potential (Ψw), thereby disrupting absorption of water by plant roots (Chutia and Borah 2012). PEG-based models of drought stress represent the method of choice in molecular biology and plant protectant studies and screening experiments (Rao and Ftz 2013).

2-3 week old barley seedlings usually have 2 emerged leaves and the beginnings of a third starting to emerge. Seedlings that reached this phase by week 3 were weighed in the same way as in the barley growth curve experiment. Osmotic stress treatment was applied by re-filling cleaned tubes with modified Hoagland’s solution with 15% PEG (8000 polyethene glycol *Sigma-aldrich,* 150g dissolved per litre of modified Hoagland’s solution) to 50 mL for every plant. Plants were returned to the controlled growth conditions and left to grow for 7 more days. Fresh and dry weight (g) were measured in roots and shoots. Dry weight was measured after plant material was placed in a paper bag and dried at 65°C for at least 7 days.

### Barley stress experiment - focus on maturation rate heterogeneity

This experiment was only carried out in the Bowman, eam8.w and eam8.k lines. Seed sterilisation, stratification, and growth conditions were the same as the barley growth curve experiment.

In this experiment, we wished to capture how heterogeneity in growth rates influences response to osmotic stress. When plants appeared to have a mass close to 0.4g, they were weighed every day. Once an individual plant reached 0.5 g±0.05g in total weight, the modified Hoagland’s solution in the tube was replaced and refilled to 50ml and the plants were left in the growing cabinet for 14 more days, disturbed only to check the solution levels, replenishing if plants had only 20 mL solution left. On the 14th day, a fresh weight was measured and a 15% PEG solution was applied as before. Wet and dry biomass of roots and shoots were measured as before after 7 days of osmotic stress treatment.

## Acknowledgements

We thank Jason Daff, Paul Scott, Harry Stevens and the rest of the University of York Horticulture team. Seeds were provided by Maria von Korff, Max Planck Institute for Plant Breeding Research, Köln, Germany.

## Competing Interests

The authors have no competing interests to declare.

## Funding

This work was supported by funding from the Biotechnology and Biological Sciences Research Council (BBSRC)—DE and SJD: BB/V006665/1. The authors also acknowledge BBSRC White Rose DTP studentships (BB/M011151/1 and BB/T007222/1) to JR (Ref.: 1792522) and EJR (Ref.: 2444228). MQ was supported by the Deutsche Forschungsgemeinschaft (DFG—514901783) through the Collaborative Research Centre 1664 “Plant Proteoform Diversity - SNP2Prot” (project A02).

## Data and resource availability

All the raw data alongside every script to generate each figure is available on: https://github.com/stressedplants/ELF3Asynchrony

## Supplemental materials

**Appendix:** Further description of the barley lines

**Fig S1:** leaf count is associated with chronological time in plants with diverse photoperiod exposures

**Fig S2:** Optimisation of germination synchronization

**Fig S3**: Barley growth curves before aligning by germination time

**Fig S4**: Early maturing barley genotypes do not display significant differences in osmotic stress response at an early time point.

## Notes

https://github.com/stressedplants/ELF3Asynchrony

### Competing Interest Statement

The authors have declared no competing interest.

### Summary of Updates

The university address of one of the authors was slightly wrong and one of the co-authors requested that we add three more relevant citations and switch our use of the word "growth" for "development" for greater scientific precision.

