## Supplemental Materials for "ELF3 controls trait heterogeneity by tuning the rate of maturation in Arabidopsis and barley"

### Supplementary materials for: “ELF3 controls trait heterogeneity by tuning the rate of maturation in Arabidopsis and barley”

All the raw data alongside every script to generate each figure is available on: https://github.com/stressedplants/ELF3Asynchrony

This supplementary material includes the following:

**Appendix:** Further description of the barley lines

**Fig S1:** leaf count is associated with chronological time in plants with diverse photoperiod exposures

**Fig S2:** Optimisation of germination synchronization

**Fig S3**: Barley growth curves before aligning by germination time

**Fig S4**: Early maturing barley genotypes do not display significant differences in osmotic stress response at an early time point.

#### **Appendix:** Further description of the barley lines

The allele for HvELF3 is located on the end of the long arm of barley chromosome 1H. B289 or eam8.k is a complex mutation involving insertions and deletions (Zakhrabekova et al., 2012). Specifically, relative to the wild type allele, it has an insertion of AGCTGCATGGCG at position 1,189 from the start of the allele followed by a deletion of 2,466 bp at position 1,189–3,656, directly followed by an inversion of bps 3,657–4,697, immediately followed by an insertion of CCGTCTCCTCCGCCTCCGCACCGTT and a deletion of 147 bp at position 4,698–4,845. These changes causes a substantial change in the gene and contribute to the loss of function in the Hvelf3 allele. B290 or eam8.w is a naturally occurring allele in the cultivar Early Russian as a single nucleotide T to C point mutation at position 2109 or in the cultivar Mona as a 4-bp deletion, both in exon 2 of the wild type allele and both result in premature stop codons causing a recessive Hvelf3 (Faure et al., 2012). Early maturity occurs due to the mutation in HvELF3 causing up-regulation of Ppd-H1 and the downstream HvFT1 under noninductive short day conditions (Faure et al., 2012). The alleles for HvLUX1 are located on the long arm of chromosome 3H below the marker ABC166 (Campoli et al., 2013), B284 or eam10.m is a nonsynonymous single nucleotide polymorphism within the exon region encoding the GARP family MYB domain, changing the wild type A nucleotide to a T in the mutant resulting in an encoded S to C amino acid change in the highly conserved SHLQKY(R/Q) motif in Hvlux1 resulting in similar phenotypes to that observed in eam8. Under short days mutation in HvLUX causes up-regulation of PpdH1 and HvPRR1 during light hours (diurnal or constant) and causes strong down regulation in HcCCA1 expression in subjective days. Unlike mutations in HvELF3 that cause compromised expression of clock oscillator and output genes other clock HvPRR genes were not as strongly affected by the Hvlux allele (Campoli et al., 2012a). The HvPHYC locus was mapped on to chromosome 5H (Pankin et al., 2014), the eam5 mutation is caused by a nonsynonymous single nucleotide mutation (T/C) in exon 1 of the HvPHYC gene. The change results in a missense substitution when the encoded F (hydrophobic phenylalanine) amino acid is replaced by a S (hydrophilic serine) within a highly conserved HHTSPRFVP (F/S)PLRYA motif in the GAF domain of phytochromes. Similar to Hvelf3 and Hvlux1 alleles the HvPHYC mutants displays photoperiod insensitivity as it interacts with Ppd-H1 and causes early maturity under long and short days, it was also found to act on the same pathway as HvELF3 and HvLUX1 disrupting circadian clock genes (Pankin et al., 2014).

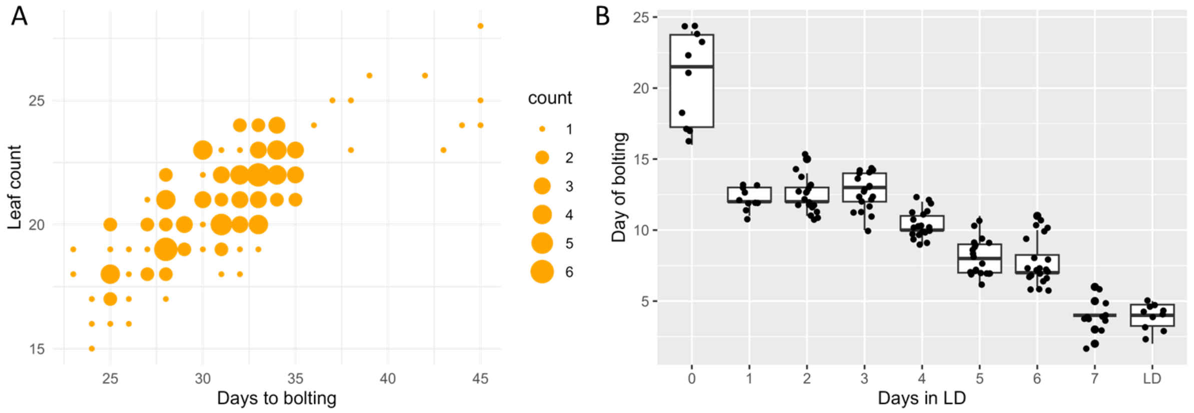


**Figure S1: Leaf count is associated with chronological time in plants with diverse photoperiod exposures.** To diversify the flowering time of Ws-2, individual plants were exposed to different durations of long day (LD, 16 hrs light/8hrs dark) treatment before being returned to short day conditions. (A) First, we compare days to bolting and the leaf count, observing a positive correlation between these two variables. (B) The boxplot illustrates how the days to bolting are negatively associated with the number of days of exposure to LD conditions. Boxplots indicate the median and interquartile range. Outliers are defined as being more than 1.5 times the interquartile range away from the median.

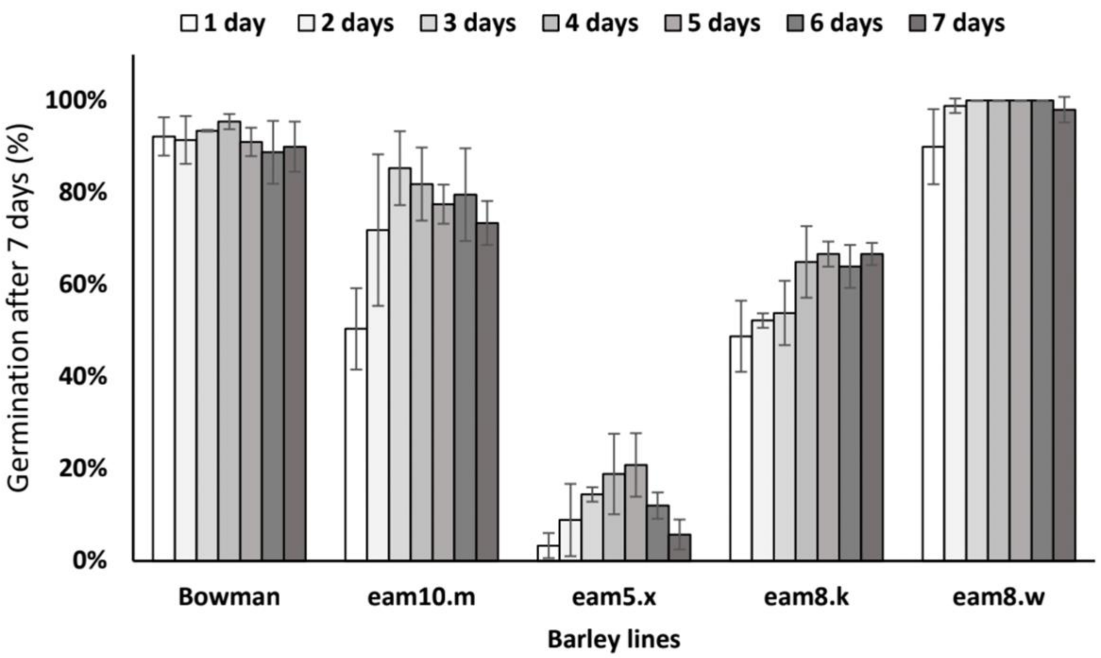


**Fig S2: Optimisation of germination.** To increase germination efficiency, we varied the duration that seeds were stratified at 4C. We observe that *eam10.m, eam5.x*, and *eam8.k* have lower germination rates than Bowman. Based on this experiment, we decided to stratify seeds for four days.

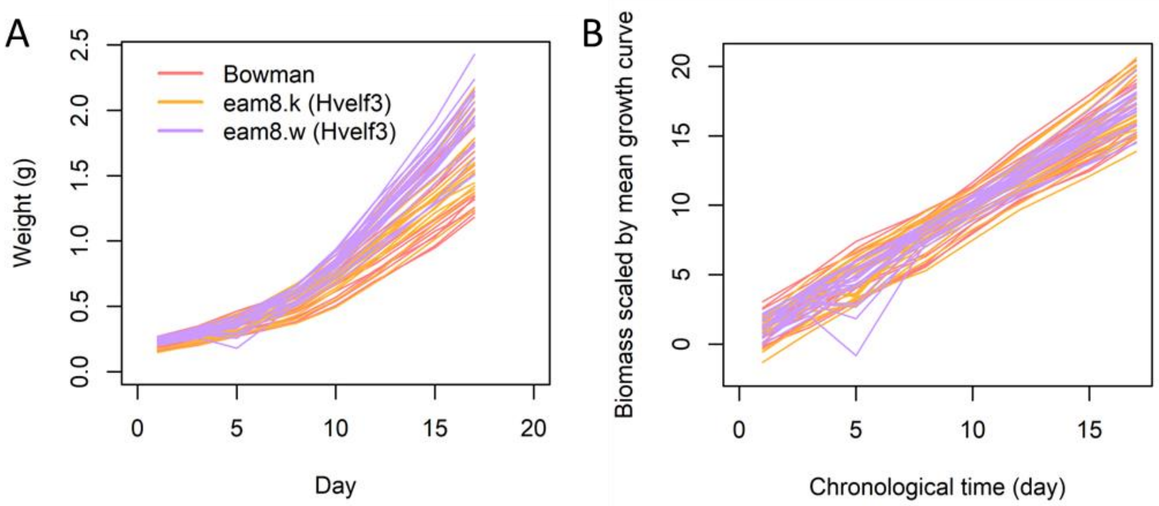


**Fig S3: Barley growth curves before aligning by germination time.** First, we show the raw growth curves, measuring weight (g) over a time series in Bowman and two variants of Hvelf3 (A). These growth curves were then aligned to the average growth curve for each genotype (B). The fitted curves were extrapolated to find their x-intercepts. Then, the day was adjusted so that the curve would intersect the x-axis on the same day. The result of this alignment is shown in **Fig 4**.

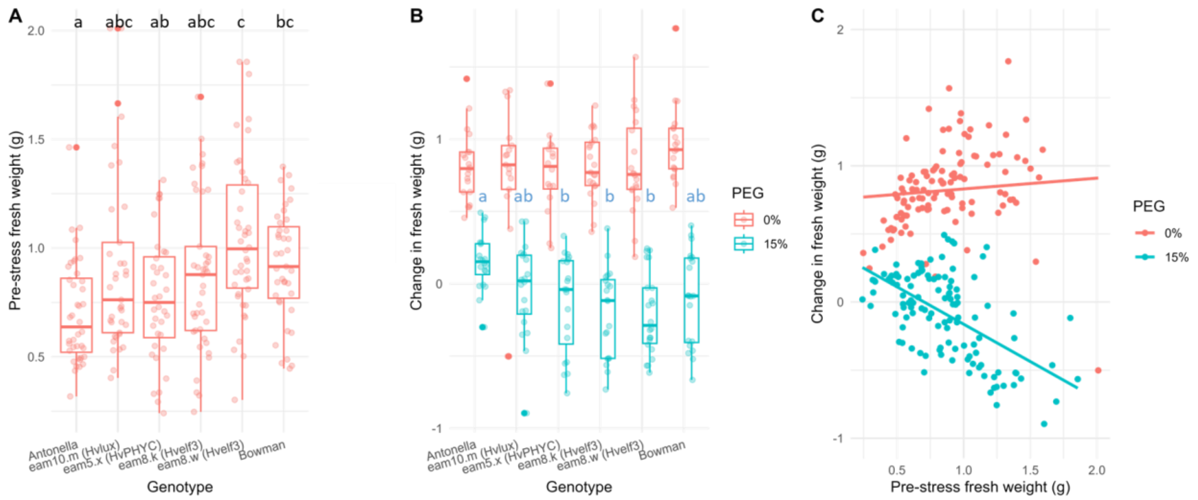


**Fig S4**: **Early maturing barley genotypes do not display significant differences in osmotic stress response at an early time point.** Several early maturing genotypes were exposed to a 15% PEG treatment to test for their water loss in response to osmotic stress. (A) It was found that there was very little variation in size at the time point in which the experiment was conducted, although Antonella had significantly lower expression than *eam8.w* and Bowman,and *eam8.w* had significantly higher expression than *eam5.x*. Boxplots show medians and interquartile ranges. Tukey HSD tests were performed at a $p<0.05$ significance level. (B) Next, we observed the change in fresh weight (g) in each of the lines under control and 15% PEG treatments, to induce osmotic stress. We only find that Antonella is significantly different from *eam5.x, eam8.k,* and *eam8.w*. Notably, none of the early maturing variants of Bowman were significantly different from Bowman. (C) However, there was a negative association between pre-stress fresh weight (g) and change in fresh weight (g). Qualitatively, the larger plants appeared more wilted, suggesting that the loss of fresh weight had a negative effect on physiology. This was even more noted in the follow-up experiment in **Fig 6**, where yellow leaves were associated with increased biomass loss. Together, this suggested to us that the size of the plants is associated with their response to osmotic stress, independently of the genotype.
